# Cellular Bioenergetics Effects of Sirolimus on the Gonads of BALB/c Mice

**DOI:** 10.1101/2023.05.15.540776

**Authors:** Hassib Narchi, Alia Al Bawardi, Pramathan Thachillath, Abdul-Kader Souid

## Abstract

**Background:** Sirolimus, an immunosuppressive drug widely used in organ transplantation, inhibits the mechanistic target of rapamycin (mTOR), decreases cellular respiration, stops cell cycle progression and causes apoptosis. It has been linked to cytotoxicity in the gonads, but the cause is unknown.

**Aims:** To understand the effects of sirolimus on the gonads by measuring cellular respiration and ATP as indicators of cellular damage, and examining histological changes.

**Material and Methods:** We used 6-8 week old male and female BALB/c mice. They received 5 µg/g sirolimus via intraperitoneal injections for 5 consecutive days per week for up to 3 weeks (treated group) or DMSO (control group). Upon euthanasia, the gonads were promptly removed while the heart was still beating, weighed, and processed for cellular respiration, ATP measurement, and histological studies. Cellular respiration was measured using a phosphorescence oxygen analyzer and ATP using a bioluminescent assay system. Histology was performed using processed tissue fragments stained with hematoxylin and eosin (H&E).

**Results:** Cellular ATP levels in testicular tissue were higher than ovarian tissue in both groups, but this was not significant (*p*=0.4 and 0.2). Although the sirolimus group had higher cellular ATP levels, no significant difference was observed in either testicular (p=0.6) or ovarian (*p*=0.9) tissue. Both DMSO (*p*=0.01) and sirolimus groups (*p*=0.008) showed lower cellular respiration in ovarian than testicular tissue. Cellular respiration was lower in sirolimus group than DMSO group in both testicular (*p*=0.6) and ovarian (*p*=0.2) tissue. Testicular cellular respiration remained unchanged up to day 11 before declining, while ovarian respiration reduced up to day 11 and remained unchanged. Testicular histopathology showed normal sperm production and normal follicular development in ovarian tissue.

**Conclusion:** The results suggest that the 5 µg/g dose of sirolimus does not cause significant short-term changes in the cellular bioenergetics of the gonads or produce noticeable histopathological changes.

## Introduction

Sirolimus, an mTOR inhibitor, is widely used in organ transplantation due to its low adverse side effects on bone marrow and kidney. It reduces cellular respiration, stops cell growth and causes cell death via apoptosis.^[1-4]^ However, it has been linked to gonadal cytotoxicity and reduced fertility in both sexes.^[5,6]^ The mechanism behind these effects is unclear, making it necessary to study sirolimus’ impact on gonadal cellular bioenergetics to fully understand its effects.

Using animal models is crucial for studying the effects of drugs on gonads, as taking samples from human gonads is unethical. Mice are an ideal model for studying cellular bioenergetics, apoptosis, and histological changes due to their suitability for pharmacological experiments, as supported by multiple studies.^[7,8]^ Measuring cellular bioenergetics with the oxygen analyzer is a well-established method in the literature.^[9, 10]^

The goal of this initial study is to establish the viability of using an animal model to investigate the impact of cytotoxic drugs, such as sirolimus, on gonadal health. The study measures cellular bioenergetics (cellular respiration and ATP) as indicators of cell damage and examines histological changes.

## Material and Methods

The study was approved by our institution’s Animal Research Ethics Committee (ERA_2016_5450).

Male and female BALB/c mice, between six and weeks of age, housed at 22°C with 60% humidity and 12-h light-dark cycles were used. They had *ad libitum* access to rodent chow and filtered water. Mice received intraperitoneal injections of sirolimus (treated groups) or DMSO (control group) for five uninterrupted days every week for 4 weeks. Sirolimus solution was prepared immediately before use by diluting the 50 µg/µL original stock in DMSO with dH_2_O to a final concentration of 0.5 µg/µL. The sirolimus dose was 5 µg/g. For controls, DMSO was diluted 100-fold with dH_2_O and injected at 10 µL/g (actual DMSO dose, 0.1 µL/g). It is worth noting that the LD_50_ of intraperitoneal sirolimus in mice is 597 µg/g (Safety Data Sheet from LC Laboratories,165 New Boston Street, Woburn, MA 01801 USA).

Weekly body weight changes (mean ± SD) in the groups of mice who received sirolimus or DMSO were measured and expressed as percentages (daily weight divided by starting weight), as the change in the body weight has been previously shown a good surrogate marker of this drug.^[11, 12]^

Immediately before each experiment, euthanasia was performed on the animals as per our institution’s animal laboratory policies. This procedure was accomplished with the use of urethane (25% w/v, 10 µL/g, administered intraperitoneally). After the animal stops moving and breathing and while the mouse’s heart is still beating, the gonads were quickly removed with a sterile scalpel and immersed in ice-cold RPMI medium saturated with 95% O_2_: 5% CO_2_. Fragments of 10 mg were collected, weighed and processed for measuring cellular respiration and ATP and for histological studies. One tissue fragment was sent for histological studies, another fragment was *immediately* immersed in ice-cold 2% trichloroacetic acid and processed for ATP determination, and another fragment was *immediately* placed in the oxygen vial for measuring the rate of cellular respiration as discussed below.

### Measurement of cellular ATP

A small organ tissue fragment (<10 mg) was immersed in 1.0 mL ice-cold 2% trichloroacetic acid (prepared daily) and rapidly homogenized by vigorous vortexing. The samples were centrifuged at -8oC at 16,873 x*g* for 10 min, and the supernatants were stored at -200C until analysis. For ATP determination, 50 µL of the cellular acid extracts were thawed on ice and neutralized with 50 µL of 100 mM Tris-acetate and 2 mM ethylenediaminetetraacetic acid (pH 7.75). The measurements used the Enliten ATP Assay System (Bioluminescence Detection Kit, Promega, Madison, WI). The luminescence reaction contained 5 µL of neutralized acid-soluble supernatant and 45 µL of luciferin/luciferase reagent. The luminescence intensity was measured at 25oC using the Glomax Luminometer (Promega, Madison, WI). ATP contents were expressed in nmol/mg protein (using the Bio-Rad Protein Assay Cat. #500-0006, Bio-Rad, Hercules, California, USA).

### Measurement of cellular respiration

The phosphorescence oxygen analyzer that measures dissolved O_2_ concentration ([O_2_]) in solution as a function of time was used to determine the rate of cellular respiration.^[9, 10]^ This method was based on the principle that O_2_ quenches the phosphorescence of a palladium (Pd) phosphor. The Pd (II) derivative of meso-tetra-(4-sulfonatophenyl)-tetrabenzoporphyrin (with an absorption maximum at 625 nm and a phosphorescence emission maximum at 800 nm) was used in this study for this purpose.

Gonadal fragments were excised and immediately placed in vials containing 1-ml phosphate-buffered saline (PBS), 3 μM Pd phosphor, and 0.5% fat-free albumin for the O_2_ measurement at 37°C. The Pd phosphor solution was prepared daily, kept on ice, and warmed to 37°C prior to use. The vials were sealed with crimp top aluminum seals. Mixing was performed with the aid of parylene-coated stirring bars. Samples were exposed to light flashes (10 per sec) from a pulsed light-emitting diode array with a peak output at 625 nm. Emitted phosphorescent light was detected by a Hamamatsu photomultiplier tube after first passing it through a wide-band interference filter centered at 800 nm. Amplified phosphorescence was digitized at 1–2 MHz using an analog/digital converter (PCI-DAS 4020/12 I/O Board) with 1 to 20 MHz outputs. The phosphorescence decay rate (1/τ) was characterized by a single exponential; I = Ae^-t^/τ, where I = Pd phosphor phosphorescence intensity. The values of 1/τ were linear with dissolved O_2_: 1/τ = 1/τo + *k*q[O2], where 1/τ = the phosphorescence decay rate in the presence of O_2_, 1/τ_o_ = the phosphorescence decay rate in the absence of O_2_, and *k*_q_ = the second-order O_2_ quenching rate constant in sec^-1^ μM^-1^.

A software package developed using Microsoft Visual Basic 6, Microsoft Access Database 2007, and Universal Library components (Universal Library for Measurements Computing Devices, http://www.mccdaq.com/daq-software/universal-library.aspx) was used for the analysis. It allowed direct reading from the PCI-DAS 4020/12 I/O Board (PCI-DAS 4020/12 I/O Board, http://www.mccdaq.com/pci-data-acquisition/PCI-DAS4020-12.aspx). The software included relational databases that stores experiments, pulses, and pulse metadata, including slopes. Pulse detection was accomplished by searching for 10 phosphorescence intensities greater than 1.0 V (by default). Peak detection was accomplished by searching for the highest 10 data points of a pulse and choosing the data point closest to the pulse decay curve from the 10 highest data points of a pulse. Depending on the sample rate, a minimum number of data points per pulse was set and used as a cutoff to remove invalid pulses with too few data points.^[10]^ The results of O_2_ consumption were expressed as µM O_2_ min^-1^ mg^-1^ tissue.

### Histopathology examination

Tissue fragments were fixed in 10% formalin and embedded in paraffin. Four-micron sections were cut and stained with hematoxylin and eosin (H&E). Additional sections were obtained on coated slides to perform immunostaining studies.

### Statistical analysis

Data was analyzed with STATA version 14, using t-tests and a Gaussian generalized linear model to observe trends. Statistical significance was determined by a 2-sided *p* value < 0.05.

## Results

### Body weight (Fig. 1)

The body weight of mice treated with sirolimus showed an initial decrease followed by a steady increase, reaching 110% at day 21. The sirolimus group consistently had a greater percentage increase in body weight than the DMSO group after the first week.

**Fig. 1.**
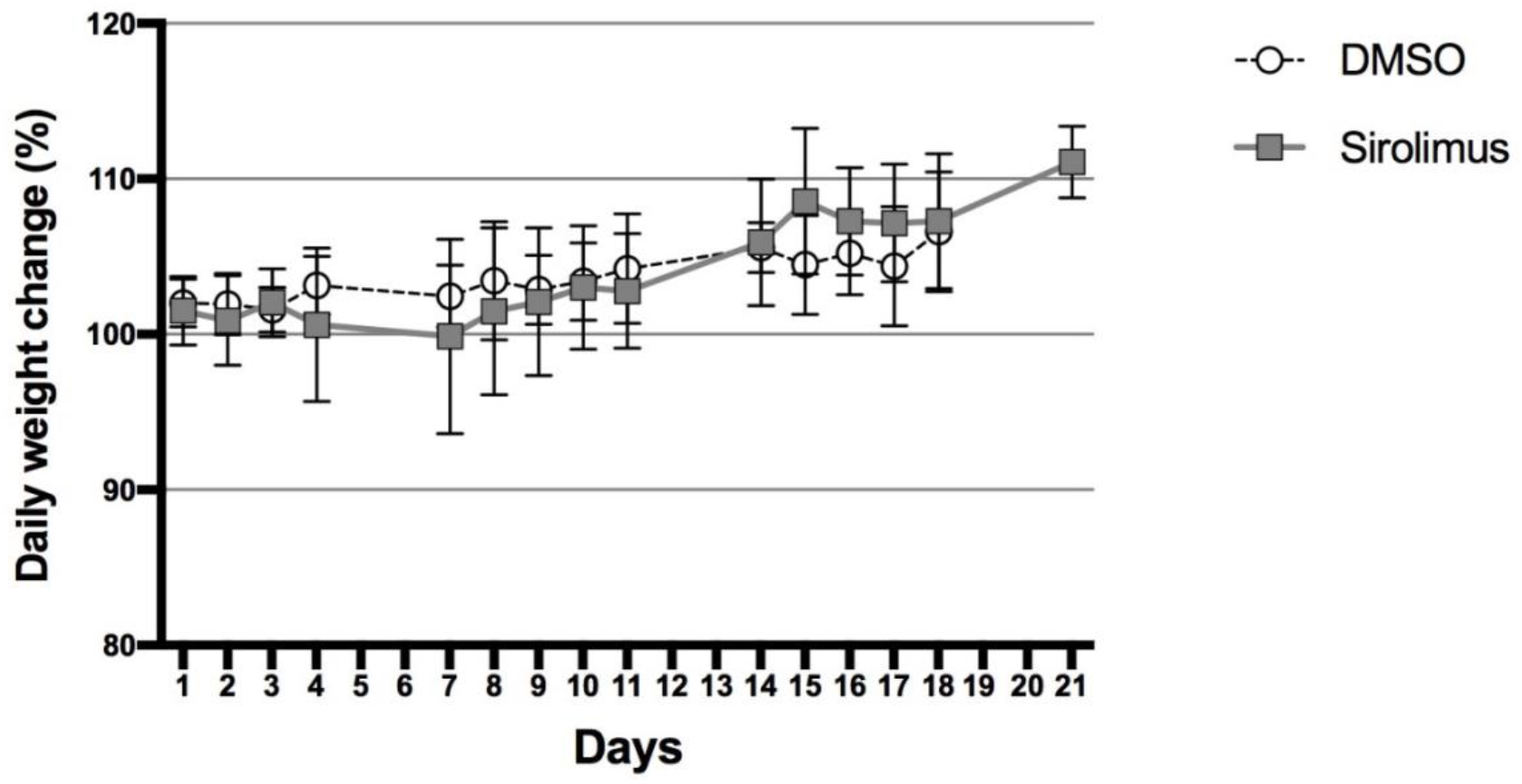
Effects of sirolimus treatment of the mouse body weight change (mean ± standard deviation) expressed as percentage of daily weight divided by starting weight.

### Cellular ATP concentration (Table 1 and Fig. 2A)

Testicular and ovarian ATP levels were higher in both DMSO and sirolimus groups compared to each other, but the difference was not statistically significant (*p*=0.4 and 0.2 for testis, and *p*=0.6 and 0.9 for ovary).

**Table 1.** Gonad cellular bioenergetics (days 4, 11, and 18). Results expressed as mean ± standard deviatio

|  | DMSO | sirolimus (5 g/g) | *p* * |
| --- | --- | --- | --- |
| *Cellular ATP (nmol/mg protein)* |  |  |  |
| Testes | 10.4 ± 7.8 (n = 5) | 8.8 ± 4.7 (n = 10) | 0.6 |
| Ovaries | 6.5 ± 7.1 (n = 5) | 6.6 ± 4.2 (n = 9) | 0.9 |
| *p* * | 0.4 | 0.2 |  |
| *Cellular Respiration (μM O<sub>2</sub> min<sup>-1</sup> mg<sup>-1</sup>)* |  |  |  |
| Testes | 0.38 ± 0.05 (n = 3) | 0.35 ± 0.06 (n = 3) | 0.6 |
| Ovaries | 0.22 ± 0.03 (n = 3) | 0.19 ± 0.02 (n = 3) | 0.2 |
| *p* * | 0.01 | 0.008 |  |

\*unpaired *t*-test

**Fig. 2.**
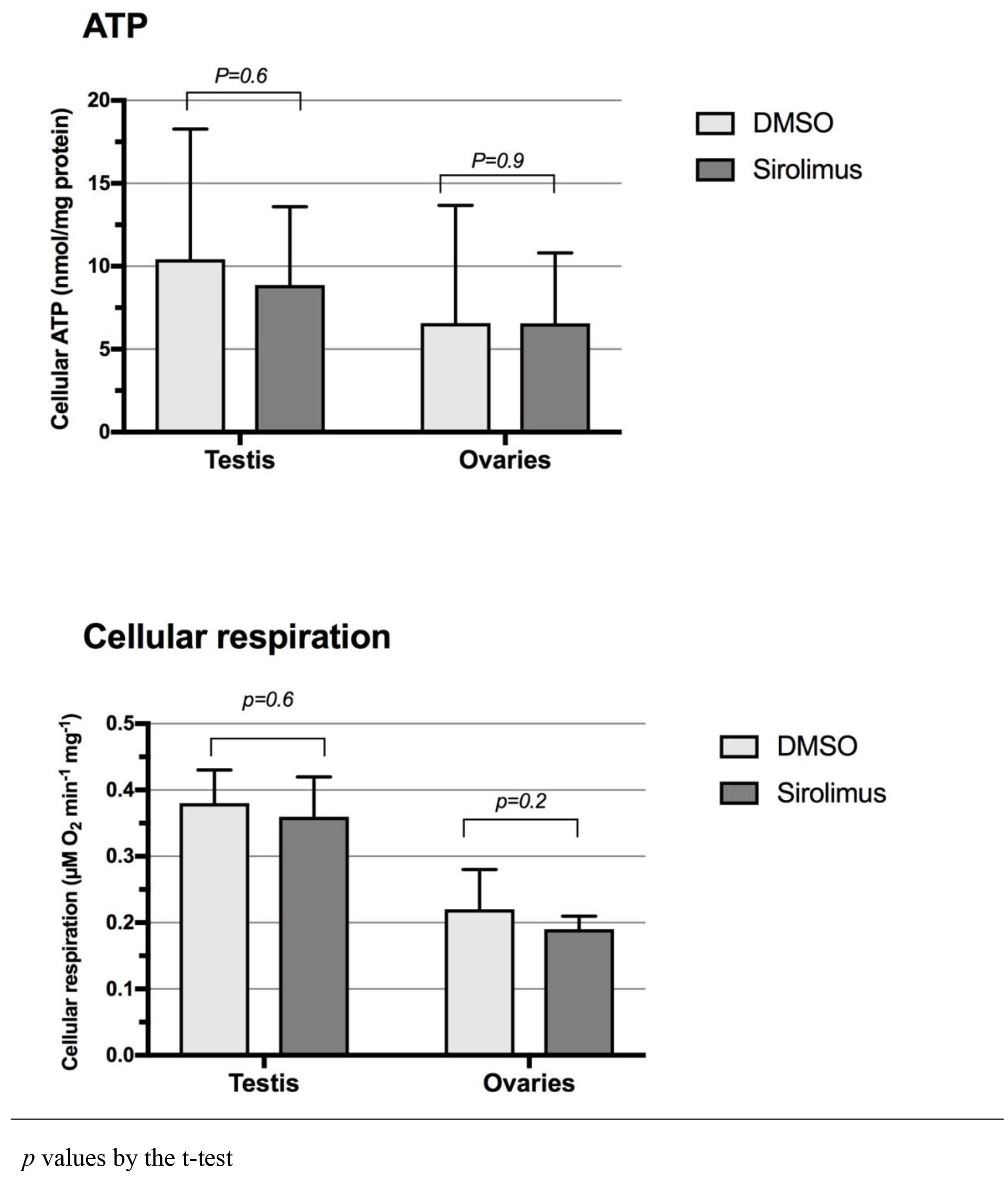
Cellular respiration (mean ± standard deviation) in the gonads (days 4, 11, and 18 after administration of sirolimus or DMSO).

After sirolimus administration, cellular ATP increased in testicles between day 4 and 11 (Fig. 3C) but decreased afterwards. No significant difference was observed in the ovarian tissue between the two groups (Fig. 3A).

**Fig. 3.**
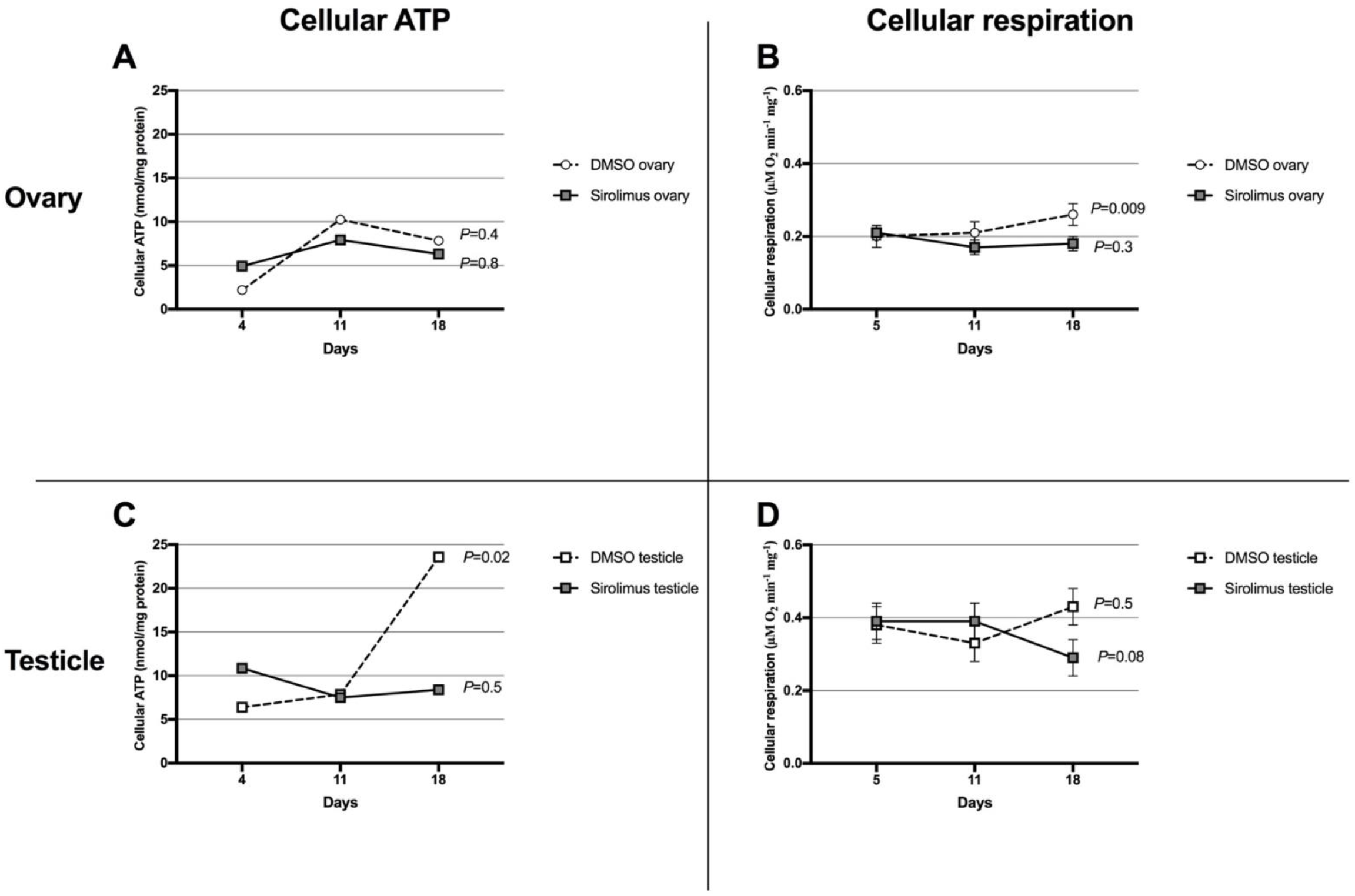
Cellular respiration trend (mean ± standard deviation) in the gonads following administration of sirolimus or DMSO

### Cellular respiration (Table 1 and Fig. 2B)

Ovarian tissue had lower cellular respiration compared to testicular tissue in both the DMSO group (*p*=0.01) and sirolimus group (*p*=0.008).

There was no significant difference in cellular respiration between sirolimus and DMSO groups in ovarian tissue (*p*=0.2). In testicular tissue, the difference in cellular respiration between sirolimus and DMSO groups was also not significant (*p*=0.6).

After sirolimus administration, cellular respiration in testicular tissue (Fig. 3D) showed no statistically significant changes (*p*=0.08) until day 11, when it decreased. Similarly, in ovarian tissue (Fig. 3B), there was a decrease in cellular respiration until day 11, but it was not statistically significant (*p*=0.3).

### Histopathogy

Histological examination of the testes revealed normal sperm production in both DMSO and sirolimus-treated mice, and the presence of secondary follicles and luteinized stromal cells in ovarian tissue indicated normal follicle development (Fig. 4).

**Fig. 4.**
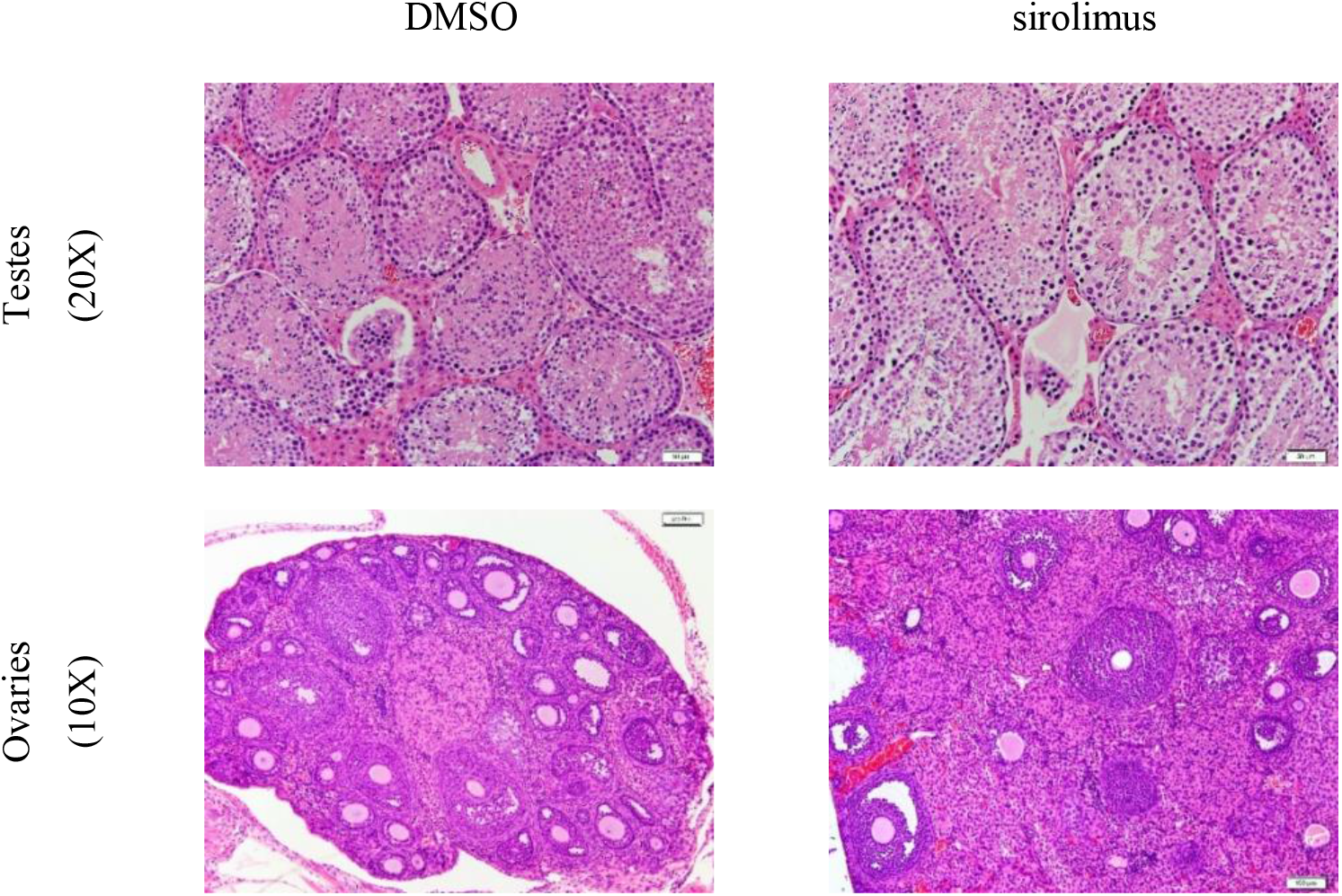
Gonads (Hematoxylin & Eosin, intermediate power). Day 4, Testes show active spermatogenesis in both DMSO and sirolimus treated mice. Day 11, Ovaries show normal follicular development, secondary follicles formation and luteinized stromal cells

## Discussion

The decrease in body weight in sirolimus-treated mice during the first week confirms drug efficacy.^[11, 12]^ The weight loss stabilized 10 days after dosing, and the animals gained weight steadily.

Cellular respiration and ATP were lower in ovarian tissue than in testicular tissue in both DMSO and sirolimus groups, indicating higher metabolism rate in testis compared to ovary even after sirolimus administration.

The administration of sirolimus did not impact cellular ATP or cellular respiration in either testicular or ovarian tissues, as the results did not show any significant changes from baseline. The cellular bioenergetics of testicular tissue remained unaffected by sirolimus, while a temporary increase in cellular ATP in ovaries was seen and quickly reversed after day 11, confirming previous reports.^[5, 13]^. Histological examination revealed no differences between the experimental groups in either ovary or testis.

The effects of mTOR on testicular function, including fertility and semen parameters, are not well understood. Known side effects of sirolimus, an mTOR inhibitor, in transplant patients are hypogonadism, reversible oligospermia, and erectile dysfunction, which are dose and duration dependent. High doses of the drug in animal models have resulted in testicular tubular atrophy. Studies have shown that sirolimus affects a crucial step in the proliferation and differentiation of spermatogonia by inhibiting the binding of c-kit to a stem cell factor through its effect on the phosphoinositide 3-kinase/mTOR pathway.^[14-16]^

This pilot study’s strength lies in establishing and demonstrating a new experimental method to examine the impact and mechanism of sirolimus on gonads’ cellular respiration and bioenergetics.

This study has limitations, such as the use of a sirolimus dose that negatively impacts body weight. Further exploration of other doses and regimens may be necessary. The role of other commonly used drugs in transplant medicine was not investigated. More detailed histopathological studies, including electron microscopy, will be conducted in future studies, as no morphological changes in gonad histopathology were observed in this pilot experiment. Possible undetectable ultrastructural changes, particularly related to the mitochondria, will be investigated in future studies, as well as the study of apoptosis. Future experiments will also consider the duration and dose dependency of sirolimus’ effects on gonadal tissues and include hormonal measurements and both quantitative and qualitative studies of sperm and oocytes.

## Conclusion

No significant short-term changes in gonads’ cellular bioenergetics or histopathology were observed with the sirolimus dose used in this pilot study. The minor decrease in ovarian bioenergetics in the first 11 days was statistically insignificant and appeared to be reversible. Additional studies are required to examine various doses of sirolimus, potential ultrastructural changes, and long-term impacts on the gonads’ cellular bioenergetics.

## Author contribution

HN and AKS designed the study, carried out the analysis, interpreted the data and drafted the manuscript. TP performed the oxygen and ATP measurements. AAB performed the histopathology examination. The manuscript was written through contributions of all authors. All authors have approved the final version of the manuscript.

## Acknowledgements

We are grateful to Mrs. Dhanya Saraswathiamma, Ms. Noora Al Mukaini and Dr. Junu Vazhappully George for their technical help in handling the mice and processing the tissue.

